# Fentanyl-Induced Diaphragmatic Discoordination during Overdose

**DOI:** 10.64898/2026.06.28.734963

**Authors:** Jaseph Soto-Perez, Gia E. Fisher, Sung Won Stephanie Wee, Brigitte Browe, Jaime Fernandez da Ponte, Yong-Hu Fang, Willard W Sharp, Alfredo J. Garcia

## Abstract

Synthetic opioids like fentanyl are a leading cause of overdose mortality. Although the hallmark of fentanyl overdose is ventilatory depression, fentanyl also induces tonic activation of skeletal musculature, including the diaphragm, which may advance progression of overdose towards death. While tonicity may further restrict diaphragmatic contractility, phase-specific dysregulation may also reflect a larger state of discoordination in respiratory control. Using urethane-anesthetized mice exposed to fentanyl, we test the hypothesis that fentanyl-induced diaphragm tonicity results from a loss of coordinated motor activity. Fentanyl produced two distinct phases: an initial phase of maximal ventilatory depression with preserved phasic activity, and a later phase characterized by unstable ventilation that partially rebounds, tonic diaphragmatic activation with loss of inspiratory phase dominance in EMG activity, and diminished bilateral diaphragmatic coordination. Carotid body denervation eliminated tonic activity and expiratory-phase EMG elevation, but it did not prevent hemi-diaphragm discoordination or ventilatory instability. Rhythmic brainstem slice recordings showed that bilateral preBötzinger complex burst-amplitude coupling was disrupted by μ-opioid receptor (MOR) agonism. Furthermore, disordered diaphragm activity was reversed by administration of the MOR antagonist, naloxone. Our findings reframe fentanyl overdose as a temporally evolving syndrome that involves distinct mechanisms to disrupt respiratory motor coordination.

**SIGNIFICANCE STATEMENT:** Respiratory depression is the hallmark of fentanyl overdose. We demonstrate that fentanyl actively reorganizes motor control of breathing, resulting in aberrant diaphragmatic tonicity and a loss of bilateral coordination. These phenomena emerge in temporal sequence and are driven by distinct central and peripheral contributions, suggesting fentanyl overdose is a complex multistage process and may require unique intervention strategies at different stages.

## INTRODUCTION

Deaths attributable to synthetic opioids such as fentanyl have risen dramatically over the past decade and now account for the majority of opioid overdose fatalities. Opioid-induced respiratory depression (OIRD) is regarded as the primary cause of death in overdose. Opioids suppress breathing by acting on μ-opioid receptors (MORs) expressed throughout the respiratory network, including the preBötzinger complex (preBötC), the kernel of inspiratory rhythmogenesis, and the Kölliker–Fuse nucleus, which shapes the transitions between respiratory phases (Smith et al., 1991; Ramirez et al., 2021; Bachmutsky et al., 2020; Varga et al., 2020). Effects of fentanyl extend beyond respiratory depression, evoking generalized skeletal muscle rigidity, most prominently in the thoracic musculature. This muscle rigidity, described clinically as wooden chest syndrome (WCS), can mechanically impede ventilation and increase the work of breathing (Torralva and Janowsky, 2019). Electromyographic (EMG) characterization indicates that this rigidity also encompasses the diaphragm, where it manifests as elevated expiratory-nadir EMG activity consistent with a failure of the muscle to fully relax during exhalation (Cavallo et al., 2023; Campbell et al., 1995)

Under normal conditions, diaphragmatic EMG activity is largely confined to the inspiratory phase (Tagliabue et al., 2024; Richter, 1982). The timing of these transitions among inspiration, post-inspiration, and expiration is governed by reciprocal interactions across distributed pontomedullary nuclei, including the Kölliker–Fuse nucleus, preBötC, the post-inspiratory complex, and the parafacial respiratory group (Dutschmann et al., 2014; Mörschel and Dutschmann, 2009). The appearance of fentanyl-driven diaphragmatic activity outside its normal inspiratory window raises the possibility that synthetic opioids disrupt phase-specific communication across premotor control centers that normally restrict diaphragmatic contraction to inspiration. The potential consequence of such loss is readily illustrated at the level of the preBötC, where bilateral premotor coordination between the left and right preBötC leads to ataxic breathing that proves lethal (Bouvier et al., 2010; Wu et al., 2017). Whether the resulting aberrant diaphragmatic activity under fentanyl reflects a loss of coordinated motor output rather than a simple intensification of central depression is poorly understood.

Fentanyl-induced muscle rigidity is attenuated by lesioning of the locus coeruleus (LC), the principal source of norepinephrine throughout the brain (Lui et al., 1989; Lui et al., 1993). Phrenic motor neurons are surrounded by noradrenergic terminals (Zhan et al., 1989) that arise in part from the LC (Bruinstroop et al., 2012; Westlund et al., 1981). Moreover, LC neurons have been proposed to be chemosensitive, responding to both hypoxia (Elam et al., 1981; Yang et al., 1997) and hypercapnia (Biancardi et al., 2008), and they receive indirect peripheral chemosensory input from the carotid bodies by way of the nucleus tractus solitarius (Lopes et al., 2016). These properties place the LC at the intersection of opioid action, motor neuromodulation, and chemoreception, and raise the possibility that the blood-gas derangements occurring during OIRD contribute to fentanyl-induced diaphragmatic rigidity (fiDR). Here we seek to define how fentanyl reshapes diaphragmatic motor output and identify contributions of central and peripheral control that drive changes in diaphragmatic activity during fentanyl-induced respiratory depression.

Using a combination of diaphragmatic and accessory-muscle EMG recordings, LC fiber photometry, carotid body denervation, and isolated preBötC slice recordings, we demonstrate that fentanyl produces temporally distinct phases of respiratory motor disruption. The initial phase is characterized by ventilatory depression with little change in phasic activity of the diaphragm, while the subsequent later phase is characterized by a partial rebound from ventilatory depression with unstable breathing, tonic diaphragmatic activation, and reduced bilateral diaphragm synchrony. Total LC GCaMP activity oscillates throughout overdose, coupling to Ve and diaphragm rigidity differentially through the evolving phases. Furthermore, LC bursting does not coincide with changes in diaphragmatic rigidity, but rather coincides with transient increases in breathing. We further show tonic diaphragmatic activation requires intact carotid body afferent input, while bilateral discoordination is independent of peripheral chemosensory drive and appears to be driven by MOR-mediated disruption in the central nervous system. These findings reframe fentanyl overdose as a multidimensional syndrome involving both central rhythm disruption and active destabilization of distributed respiratory control and coordination.

## METHODS

### Animals

All animal protocols followed National Institutes of Health guidelines for the appropriate care and use of animals and were approved by the Institutional Animal Care and Use Committee at The University of Chicago. Experiments were performed in the light cycle. For fiber photometry studies, we used a transgenic mouse line where Cre-recombinase expression is driven by the norepinephrine (NE) transporter promoter and has high selectivity for NE cells in the Locus Coeruleus (LC) with low ectopic expression (slc6a2, mouse line: Net-Cre1, Wagatsuma et al., 2018). Adult mice of both sexes (>9 weeks, 22-41g) on a Net-Cre background were used for simultaneous plethysmography, EMG, and fiber photometry studies. Mice were bred at the University of Chicago animal facilities. Mice used in this study were group housed with the same sex in AAALAC-approved facilities following a 12/12 h light/dark cycle and were given ad libitum access to food and water.

#### Stereotaxic Surgical Procedures

All viral injections were done with a stereotaxic surgical device (Kopf Instruments) to position injections and implants correctly and sterile tools were used throughout to maintain a sterile surgical environment. Isoflurane (2-3% induction, 1-1.5% maintenance) was used to anesthetize the mice. Head hair was shaved using a trimmer, and the mice were head-fixed in the stereotaxic system. Sterile drapes were placed over a heating pad to maintain homeostatic body temperature, and ophthalmic lubricant was placed over the eyes. Betadine was applied on the scalp prior to any incision or injection. Bupivacaine (1 mg/kg, s.c.) was then injected into the scalp. Sterile tools were used to make an incision at the midline of the scalp. A micromotor drill attached to the stereotaxic apparatus was used to drill two small holes into opposite sides of the skull where two small screws were implanted. At the end of the surgery, mice were injected with Meloxicam (5 mg/kg, s.c.) and saline (0.5 mL) and returned to their cage. The cage was then placed on a heating tray until the mice recovered and regained normal mobility (about 1-3 days).

#### Carotid Body Denervations

Bilateral carotid body denervation (CBD) was performed as previously described (Peng et al., 2026). Briefly, mice were anesthetized with 3% isoflurane delivered in 100% oxygen and maintained at 1.5% isoflurane throughout the procedure. Following induction, an endotracheal tube was inserted, and mice were mechanically ventilated (10 mL/kg tidal volume, 110 breaths/min, and 2 cmH2O PEEP) for the duration of the surgery. Body temperature was monitored continuously and maintained at 37°C using a feedback-controlled heating lamp. With the mouse positioned supine, a midline cervical incision was made, and the underlying musculature was bluntly dissected to expose the carotid bifurcation bilaterally. The carotid sinus nerve was carefully isolated and sectioned taking care to preserve the integrity of the surrounding vasculature. The incision was closed with interrupted sutures. Mice were removed from anesthesia, extubated upon return of spontaneous breathing, and monitored during recovery. Sham-operated animals underwent identical surgical exposure of the carotid bifurcation bilaterally without nerve sectioning.

#### Viral injections and cannula implants for fiber photometry

A Hamilton syringe was attached to the stereotaxic apparatus for viral injections and slowly lowered to the Locus Coeruleus. A syringe pump was used to control the viral injection volume to 300 nL per injection. The needle was held in place for a total of 10 minutes to allow for sufficient viral infusion and was then slowly removed. To monitor the activity of the noradrenergic cells of the LC, we performed stereotaxic injections of a Cre-dependent virus (pGP-AAC-CAG-FLEX-GCaMP8s-WPRE, Addgene: 162380) that genetically encodes the Ca2+ indicator, GCaMP8 into the LC of Net^Cre/+^ mice. Fiber optic cannulas (MFC_400/430-0.66_6mm_MF1.25_FL, Doric) were implanted using a cannula holder attached to the stereotaxic system. Dental cement was applied to the screws, skull, and cannula to form a head cap that holds and protects the fiber optic cannula. Experimenters waited at least 3 weeks for the virus to fully express before checking the fluorescent signal. LC coordinates relative to Bregma for cannula placement were AP: -5.4, ML: 0.85, DV: -3.75. Two DV viral injections were performed dorsal and ventral to the coordinates - 3.9 and -3.6. DV cannula placement was at -3.75 (verified using Paxinos and Franklin’s Mouse Brain Atlas, 2001).

#### Coupled Electromyography and Spirometry

Mice were anesthetized with Urethane (1.5g/kg) and placed on a heating pad for ∼10min prior to surgery, maintaining body temperature at ∼37°C. An incision in the pectoral muscle was made and a skin pocket spanning from the pectoral to the external oblique was formed using blunt dissection scissors. The skin was resected, and muscles were bathed in saline solution. EMG leads were placed unilaterally or bilaterally in the diaphragm underneath the costal cartilage of ribs. In a subset of animals, abdominal and diaphragm muscle activity was measured concurrently. In this subset, a skin pocket exposing the abdominal wall was made and EMG leads were placed in the rectus abdominalis. Raw EMG signal was amplified 10,000X using a GRASS P55 amplifier. A custom-built helmet (5mL volume) connected to a variable pressure sensor was used to measure ventilation. A constant flow of 150-200mL was used to ensure no buildup of CO_2_ inside the helmet. Pressure deflections were amplified using a Buxco Max 2 amplifier. Both EMG and pressure signals were digitized using an Axon Instruments Minidigi 1B and recorded on Axoscope software.

Data was loaded into MATLAB using the abf2load function. Individual breaths were detected from the plethysmography signal using the find peaks algorithm with user-defined minimum peak prominence and height thresholds. Valleys flanking each peak were similarly detected on the inverted signal. Peaks exceeding a user-defined maximum amplitude ceiling were rejected. Each breath cycle was defined as the interval between two consecutive valleys. Each breath cycle was further subdivided into three physiological phases. The early expiratory phase (E1) was defined as the interval from the onset valley to the first upward zero-crossing of the plethysmography signal. The inspiratory phase (I1) spanned from that zero-crossing until the signal subsequently fell back below zero, subject to a user-defined minimum amplitude gate to exclude subthreshold deflections. The late expiratory phase (E2) was defined as the interval from the end of I1 to the subsequent valley.

EMG signal was rectified and smoothed using a moving average filter. Cardiac events were identified as rectified EMG activity exceeding a user-defined amplitude threshold. Each detected noise epoch was replaced by the immediately preceding segment of equal duration, preserving signal continuity. The resulting cleaned EMG signal was then segmented according to the E1, I1, and E2 phase boundaries identified from the plethysmography signal. For each breath cycle and each phase, time-normalized integrated EMG activity was computed as the trapezoidal integral of the cleaned EMG divided by the phase duration. All per-breath metrics, phase timing, waveform data, and time-normalized EMG values were exported as comma-separated value files for subsequent statistical analysis. Minute ventilation (Ve) was computed for each breath as the product of tidal volume and instantaneous breathing frequency. Analysis windows (20-30sec) were then specified for each epoch of interest (e.g., baseline, fentanyl, naloxone). The mean value and coefficient of variation (CV%, calculated as 100 × SD/mean) for Frequency, Tidal Volume, Ve, and minimum diaphragm EMG were computed from the defined windows and exported for subsequent statistical comparisons.

#### Fiber Photometry

To concurrently measure LC activity alongside flow and diaphragm EMG, mice were anesthetized and prepared as mentioned above. Fiber photometry was conducted using the TDT RZ10X system with a lock-in amplifier for signal demodulation, in conjunction with a Doric 6-port fluorescent mini cube (FMC6_IE(400-410)_E1(460-490)_F1(500-540)_E2(555-570)_F2(580-680)_S, Doric). The mini cube was equipped with 415 nm (isosbestic control) and 465 nm (GCaMP) excitation for our fluorophores of interest. 465 nm (GCaMP) was modulated at 330 Hz and 415 nm (isosbestic control) was modulated at 210 Hz. The signal was sampled at 1017.3 Hz. Output power levels were matched between isosbestic and GCaMP signals and were kept consistent. TDT Synapse software was utilized to record and process fluorescent signals, track power levels, and simultaneously capture breathing and electromyography signals. Photometry sessions lasted 45 minutes on average, including baseline, fentanyl and naloxone exposures.

Fiber photometry, breathing, and EMG signals acquired through Synapse software were preprocessed in MATLAB using the TDTbin2mat function. Signals were uniformly processed using windowed smoothing for down-sampling and noise reduction. For the selected interval, the GCaMP (465 nm), isosbestic (415 nm), plethysmograph, and EMG signals are extracted and smoothed using moving averages with signal-specific window sizes (10 samples for fluorescence, 20 for EMG), effectively reducing high-frequency noise while preserving slower dynamics. Raw GCaMP data was corrected for motion artifact and baseline fluctuations by fitting the isosbestic wavelength (415nm) signal to the GCaMP wavelength (465nm) signal with the polyfit function. To determine ΔF/F, the difference between the 415 fit and 465 signal was then divided by the 415 fit (ΔF/F (=[F465-Ffit415]/ Ffit415)). A Z-score of ΔF/F was used to compare across conditions. For both fentanyl and naloxone conditions, we used the mean and standard deviation from the baseline session (Urethane anesthesia alone) to calculate the Z-score. The find peaks function was used to locate LC fiber photometry events. The threshold for an event to be counted was set at a Z-score of 2. This is to ensure with 95% confidence that the events being counted were significantly different from the mean fluorescent signal. The findpeaks function was also used to identify breaths and RR, TV, Ve and minimum diaphragm activity was computed as described above and measured inside and outside of detected LC events across conditions.

#### Immunohistochemistry

Isoflurane anesthetized mice underwent transcardial perfusion with PBS followed by 4% PFA. Brains were harvested and kept on fixative overnight then transferred to 30% sucrose until further processing. 50uL slices containing the LC were collected using a vibratome. Slices were washed with PBS, blocked with 2% donkey serum and 0.2% triton X. Slices were then stained with rabbit anti-GFP (1꞉500; RRID:AB_2307355) and sheep anti-TH (1꞉400; RRID:AB_561880) primary antibodies in blocking solution overnight followed by secondary antibodies (Donkey anti-rabbit 488: 1:500, RRID: AB_2535792 ; Donkey anti-sheep Cy3, RRID: AB_2340727) staining for 2 hours. Slices were mounted on slides with Prolong Gold with dapi and cured at 4°C for 24 hours. Images were acquired at 20X magnification using an AX10 fluorescent microscope.

#### Brain Slice Electrophysiology

Pups (pnd 7-11) were euthanized by rapid decapitation, and brainstems were subsequently dissected, isolated, and placed into artificial cerebrospinal fluid (aCSF) (composition in mM: 118 NaCl, 25 NaHCO_3_, 1 NaH_2_PO_4_, 1 MgCl_2_, 3 KCl, 30 Glucose, 1.5 CaCl_2_, pH = 7.4) equilibrated with 95% O_2_, 5% CO_2_ at ∼4°C. The brainstem was glued to an agar block (dorsal face to agar) with the rostral face upward and submerged in aCSF equilibrated with carbogen. Serial cuts were made through the brainstem until anatomical landmarks such as the narrowing of the fourth ventricle and the hypoglossal axons appeared. The preBötC was isolated in a single transverse brainstem slice (thickness: 650 μm). The slice was transferred into the recording chamber (∼6 mL volume), where it was continuously perfused with recirculating aCSF (flow rate: 12–15 mL/min, ∼30°C–34°C). Extracellular KCl was raised to 8 mM, and the spontaneous rhythm was allowed to stabilize before the start of every recording. Electrophysiology recordings were made in Clampex or Axoscope within the pClamp software package (Molecular Devices, San Jose, CA). Pipettes were positioned bilaterally over the ventral respiratory column containing the preBötC. Extracellular population activity was recorded with glass suction pipettes filled with aCSF (Garcia et al., 2016, Browe et al., 2023). The recorded signal was sampled at 5 kHz, amplified 10,000 X, with a lowpass filter of 10 kHz using an A-M instruments (A-M Systems, Sequim, WA, USA) extracellular amplifier. The signal was then rectified and integrated. Recordings were stored on a computer for posthoc analysis. Recordings of preBötC activity were made under baseline conditions (95% O_2_ 5% CO_2_, 600 s) for 10 minutes followed with the addition of 60nM of the mu opioid receptor (MOR) agonist, [D-Ala2, NMe-Phe4, Gly-ol5]-enkephalin (DAMGO) for 15 minutes. Bursts were considered corresponding if the initial start time of bursts were within 250ms of each other. The analysis window was taken at the end of each baseline or DAMGO phase (phase duration: baseline = 600 s, DAMGO = 900s). The burst amplitude ratio for each inspiratory event (defined by a network burst in the dominant preBötC, as determined by lowest value of amplitude irregularity) was calculated as the ratio of the dominant preBötC burst area to corresponding contralateral preBötC burst area, previously described as I/O ratio in Garcia et al., 2016 and Browe et al., 2023.

### Statistics

All statistical analyses were performed using GraphPad Prism, with data expressed as means and significance set at p < 0.05. Two-condition within-subject comparisons were assessed using paired t-tests. Comparisons across three or more repeated conditions within the same animals were evaluated using repeated-measures one-way ANOVA with Tukey’s multiple comparisons test. Phase-specific diaphragmatic activity (inspiratory versus expiratory ∫Dia EMG) and the resulting I/E ratio across conditions were analyzed by mixed-effects analysis with Tukey’s multiple comparisons. Two-factor designs were assessed using two-way ANOVA or Mixed effects analysis with either Fisher’s LSD post hoc test or Tukey’s multiple comparisons. Statistical significance is denoted by asterisks indicating differences between conditions (and, where noted, hashtags indicating differences from baseline), with one symbol corresponding to p < 0.05, two to p < 0.01, three to p < 0.001, and four to p < 0.0001.

### Code Availability

All custom MATLAB scripts used for signal preprocessing, event detection, and data analysis will be made available upon request.

## RESULTS

### Effects of fentanyl on ventilation and diaphragmatic function

Systemic fentanyl administration reliably suppressed breathing by ∼40% (Fig. 1). Concurrent with ventilatory depression (Fig 1A *top*; Fig 1B top), fentanyl (0.7mg/kg i.p.) induced tonic activation of the diaphragm in most subjects (∼75%, 11 out of 15 mice), as evidenced by elevated minimum EMG activity (Fig. 1A *bottom*, Fig 1B bottom). Following fentanyl administration, Ve declined progressively until reaching a plateau of maximal depression (time to peak Ve depression: 156 +/-18 sec after fentanyl administration). During this early phase, where maximal Ve depression occurred, minimum diaphragmatic EMG (min ∫Dia) activity showed a slight increase of ∼20%. As fentanyl exposure progressed, min ∫Dia significantly increased to over 200% on average. fiDR consistently occurred after the peak depression of Ve (time to peak fiDR: 355 +/- 47 seconds after fentanyl administration). Peak fiDR was consistently accompanied by a rebound in ventilation. Examination of the mean waveforms during baseline conditions and in the two phases of fentanyl suggested that the fentanyl-disrupted breathing patterns were present during both early and late stages (Fig 1C), which was linked with increased variability in respiratory frequency (CV%: Early = 13.67 +/- 2.3 ; Late = 26.59 +/- 3.56) and tidal volume (CV%: Early = 12.24 +/- 1.73 ; Late = 16.84 +/- 1.79) (Fig 1D). Together, these data suggest that fentanyl overdose leads to temporally distinct phases of ventilatory dysfunction, initially characterized by maximal ventilatory depression. Although ventilation partially rebounds during the later phase, breathing remains irregular, and aberrant diaphragmatic activity emerges.

**Figure 1.**
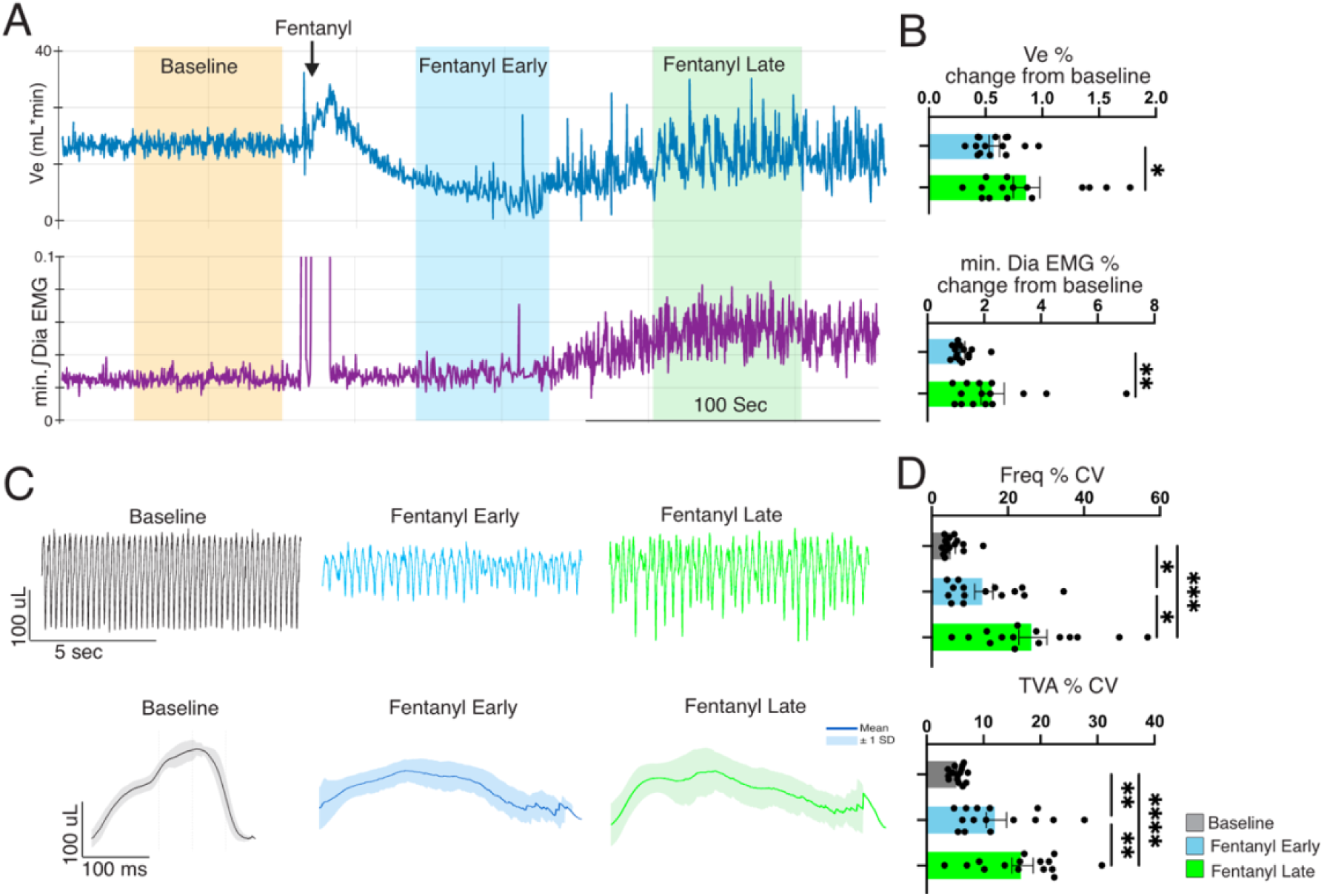
Temporal dynamics of fentanyl-induced diaphragmatic rigidity. A) Representative traces of Ve (top) and minimum integrated diaphragmatic activity (bottom) show temporal evolution of OIRD and diaphragmatic tonicity following fentanyl administration (0.7mg/kg). B) Summary data (n = 15) shows Ve rebounds (top, p = 0.021) as min ∫Dia EMG increases (bottom, p = 0.005) (Means are compared using paired t-test). C) Raw traces of respiratory activity (top) and the average breath waveform (bottom) under baseline (black) and during the early (middle) and late (right) fentanyl exposure show ventilatory instability under fentanyl worsens after peak OIRD. D) Respiratory variability was evaluated using the coefficient of variance. Summary data (n = 15) shows respiratory variability in Frequency (top, p = 0.014) and Tidal Volume (bottom, p = 0.004) progressively worsens from the early to late periods following fentanyl administration (RM One-Way ANOVA with Tukey’s multiple comparison). Asterisk (*) indicates the difference between groups; One symbol = p < 0.05, two symbols = p < 0.01, three symbols = p < 0.001, four symbols = p < 0.0001.

### fiDR coincides with disrupted phasic diaphragmatic activity

Notably, the pattern of diaphragmatic activation was not uniform across the respiratory cycle. Sectioning out breath phases into inspiration and expiration (Fig. 2A) shows fentanyl preferentially augmented EMG activity during the expiratory phase (Fig. 2Bii, p < 0.0001), a period during which the diaphragm is normally at its lowest. On the other hand, fentanyl had no effect on integrated diaphragm EMG activity in the inspiratory phase (Fig 2Bi, p = 0.68). Under normal conditions, inspiratory diaphragmatic EMG dominates throughout a respiratory cycle. However, we observe a fentanyl dependent drop in the ratio of inspiratory to expiratory EMG activity (Fig 2Biii, p < 0.0001) indicating a loss of preferential phasing of diaphragmatic activity. Tonic expiratory diaphragmatic activation was fully reversed by naloxone (2mg/kg, i.p., Fig. 2Bii, p = 0.0008), confirming that these effects are mediated through MOR signaling. Taken together, these data suggest disordered breathing in our model is driven by a loss of coordinated, inspiration dominant, motor output from the diaphragm.

**Figure 2.**
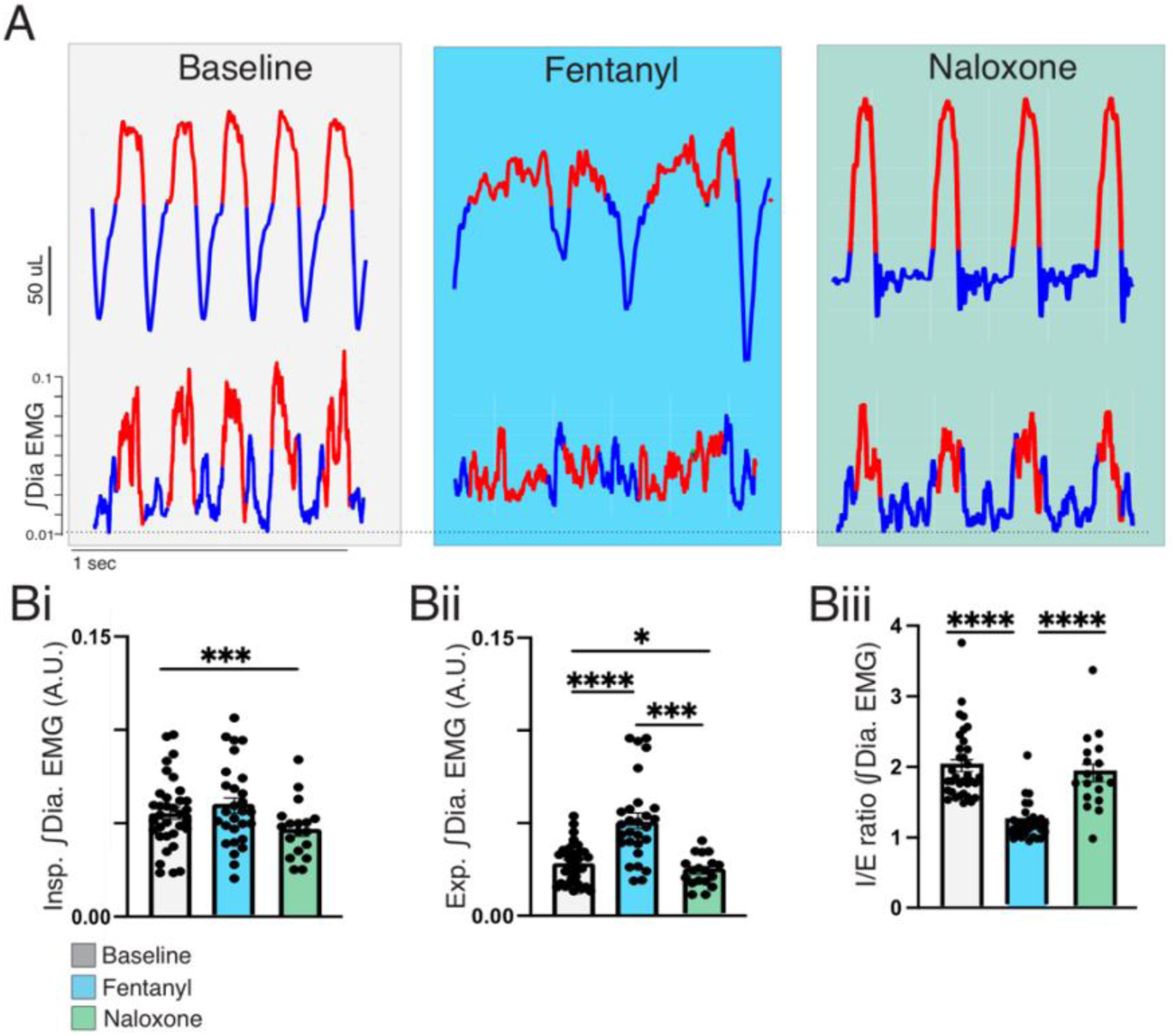
Fentanyl-induced aberrant diaphragmatic activity preferentially in the expiratory phase. A) Representative traces of ventilatory flow (top) and integrated EMG (bottom) show fentanyl (middle) depresses ventilation and increases tonic diaphragm activity. Naloxone (right, 2mg/kg, i.p.) rescued respiratory depression and tonic diaphragm activity. Tonic ∫Dia EMG was not uniformly distributed across the respiratory cycle, as dividing respiratory phases into inspiration and expiration shows elevated ∫Dia EMG found during expiration (blue), but not inspiration (red). B) Summary data (n = 34) shows, compared to baseline, ∫Dia EMG activity following fentanyl is elevated during expiration (Bii, p < 0.0001) but not different during inspiration (Bi; p = 0.68) resulting in a decrease in Inspiration/Expiration ratio (Biii, p < 0.0001). Mixed Effects Analysis with Tukey’s multiple comparisons. Asterisk (*) indicates the difference between conditions;. One symbol = p < 0.05, two symbols = p < 0.01, three symbols = p < 0.001, four symbols = p < 0.0001.

To evaluate whether fentanyl-induced changes to phase-specific activity extended to musculature beyond the diaphragm, we simultaneously recorded from the diaphragm and the rectus abdominis under baseline and fentanyl conditions. The rectus abdominis is inactive under baseline conditions and can thus serve as a tonic control of accessory musculature. While fentanyl elevated diaphragmatic EMG activity during the expiratory phase (Supplemental Fig 1Ai-ii), the rectus abdominis activity was no different from baseline in either inspiratory or expiratory phases (Supplemental Fig. 1Aiii). Muscle activity from the rectus abdominalis was confirmed under severe hypoxic challenge (5% FiO_2_). This highlights divergent susceptibility in respiratory musculature to discoordination under fentanyl and confirms that the aberrant expiratory-phase signal recorded from the diaphragm is not an artifact of crosstalk from accessory respiratory muscles.

Given tonicity observed in the diaphragm was not originating from peripheral musculature (i.e. abdominal muscles), it is possible fentanyl-induced tonicity of diaphragmatic activity may be a consequence of aberrant inputs from premotor to the phrenic motor pools. To assess whether diaphragmatic tonicity reflected dysregulated coordination of premotor inspiratory activity, we recorded bilaterally from the left and right preBötzinger complex (preBötC) using the rhythmic medullary slice preparation. Under normal conditions, bilateral preBötC activity generates phase-locked bursts which merge onto motor nuclei to produce inspiratory drive, thus making it an ideal premotor area to assess respiratory coordination. Bath application of DAMGO (60nM) reduces the near one-to-one burst amplitude relationship between the left and right preBötC (Fig. 3B, p = 0.009), supporting the notion that opioid modulation leads to the disruption of bilateral coordination at the level of the inspiratory rhythm generator. To determine whether bilateral premotor disruption translated into hemi-diaphragmatic function in vivo, EMG activity was measured from the left and right sides of the diaphragm in a separate cohort of animals. Polar plots of peak EMG activity show, under baseline conditions, diaphragmatic contractions from the left and right diaphragm are synchronized (Fig 3C). In contrast, fentanyl induced tonic EMG activity in both hemi-diaphragms (Fig 3E; Left, p = 0.011; Right, p = 0.0039) that reduced the coordination of hemi-diaphragms as indicated by a reduction in the phase-locking value (Fig. 3D, p = 0.017) which is further indicative of a loss of bilateral motor coordination.

**Figure 3.**
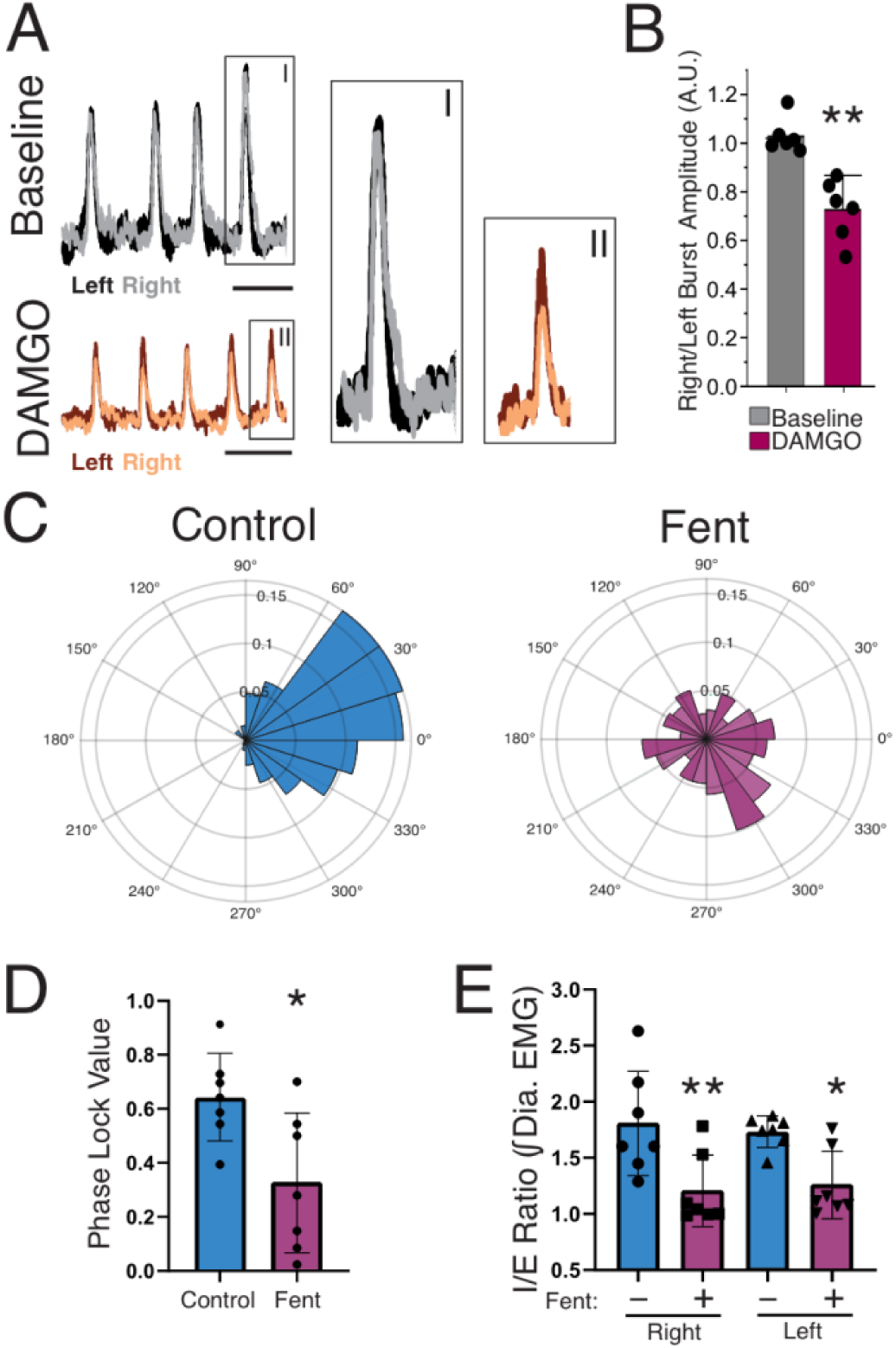
Fentanyl leads to a loss of coordinated bilateral diaphragmatic activity. A) Left, Representative traces of bilateral preBötC activity under baseline and DAMGO. Scale bar = 5 seconds. B) Summary data (n = 6) shows a shift in burst amplitude relationship (left, p = 0.009; paired t-test) from baseline (gray) to DAMGO (60nM, purple). C) Polar plots of maximum Left-Right diaphragmatic EMG activity show loss of bilateral diaphragm coordination under fentanyl conditions. D) Summary Data (n = 7) of left-right phase lock values under baseline (blue) and fentanyl (green) shows a reduction in hemidiaphragm coordination (p = 0.017; paired t-test). E) Loss of phase specific diaphragm activity was observed in both left and right hemidiaphragm as shown by summary data of I/E ratio from (Left, p = 0.011; Right, p = 0.0039; two-way ANOVA with Fishers LSD test). Asterisk (*) indicates the difference between conditions;. One symbol = p < 0.05, two symbols = p < 0.01, three symbols = p < 0.001, four symbols = p < 0.0001.

### Relationship between LC activity and fiDR

The LC is a modulator of premotor respiratory control (de Carvalho et al., 2017; Biancardi et al., 2008; Yackle et al., 2017), expresses MOR, and is involved in mediating fentanyl-induced muscle rigidity (Lui et al 1989, 1993). Therefore, the LC is a strong candidate to contribute to aberrant diaphragmatic activity under fentanyl. Despite this, no characterization of LC activity during this phenomenon and following its reversal with naloxone exists. To address this, we next examined the relationship between LC activity and fiDR using fiber photometry to track GCaMP activity from LC neurons (Fig 4A, Fig 4Bi top) while simultaneously measuring ventilation (Fig 4Bi middle) and diaphragmatic EMG (Fig 4Bi bottom). Fentanyl induced slow, stereotypical shifts in GCaMP fluorescence that initially decreased and later rebounded (Fig 4Bi top, Bii, p = 0.026). While LC dF/F was initially reduced in concurrence with the depression of Ve, both dF/F and Ve progressively recovered (Fig 4Biii, p = 0.0079). On the other hand, elevated diaphragmatic rigidity coincided with rebounding dF/F, not with its initial depression (Fig 4Biv). Close examination of the LC GCaMP activity also revealed discrete bursts of LC activity evident throughout each experimental phase (Fig 4Ci). During baseline, these transient LC events did not coincide with changes in either Ve or minimum diaphragmatic EMG. However, with fentanyl, the rate of these bursts increased (Fig 4Cii, p = 0.023) and Ve during these events was transiently stimulated (Fig. 4Ciii, p = 0.0003) while minimum diaphragmatic EMG activity was unchanged (Fig. 4Civ, p = 0.13). Following fentanyl, naloxone did not affect total LC dF/F (Supplemental Figure 2, Bi) or reduce the number of GCamp burst events (Supplemental Figure 2, Ci, p = 0.096). Furthermore, naloxone reversed Ve stimulation by LC burst (Supplemental Figure 2, Cii, p = 0.038) while preserving the relationship between LC bursts and minimum ∫Dia EMG activity (Supplemental Figure 2, Ciii, p = 0.15).

**Figure 4.**
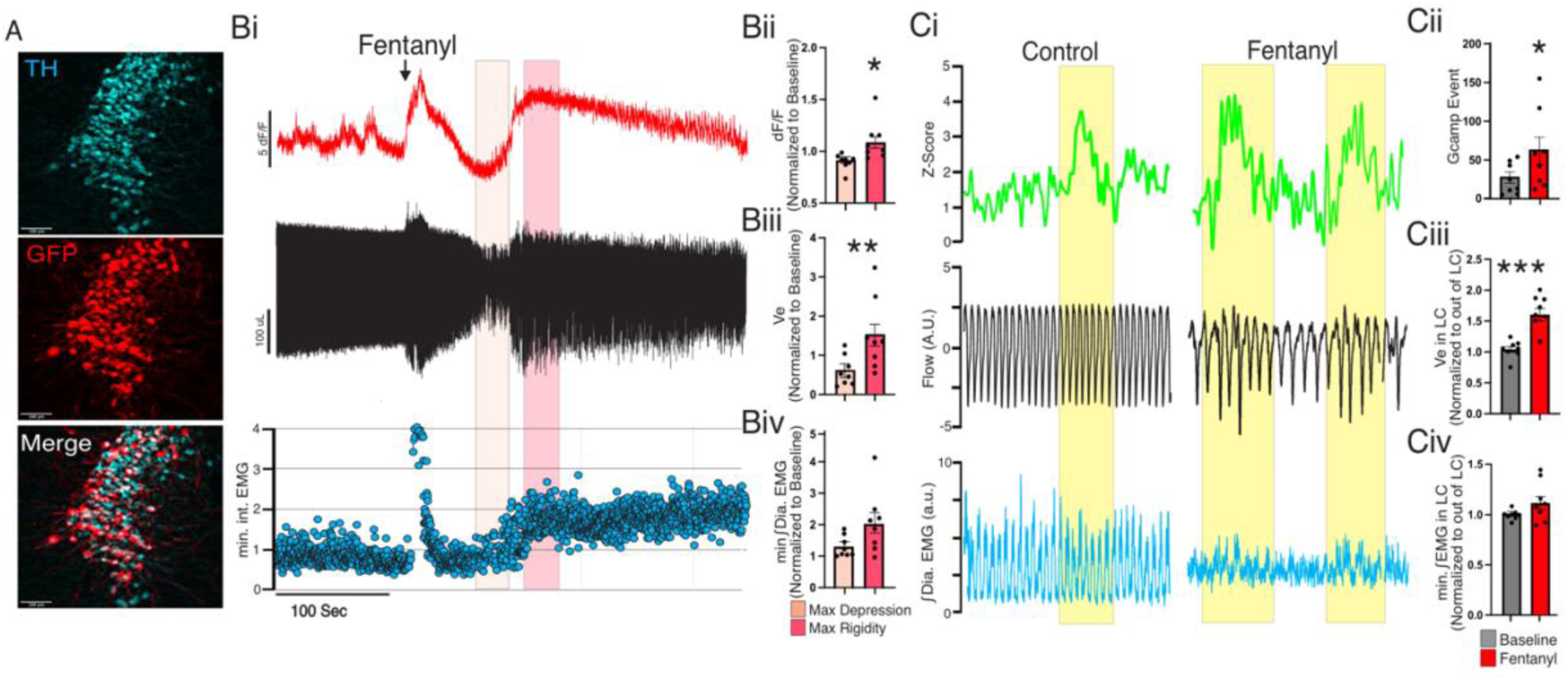
Locus Coeruleus engages in multiple activity modalities which couple differentially to diaphragm motor output and ventilation. A) Immunohistochemistry of the LC region from a Net^Cre/+^ mouse following transfection with AAV-FLEX-GCaMP8 shows Co-Expression of GFP (red) in TH+ (cyan) expressing cells. Bi) Representative trace of dF/F (Top), plethysmography (middle), and minimum Diaphragm EMG under baseline and fentanyl. Shaded areas represent period of maximum respiratory depression and rigidity onset. Bii) Summary data (n = 8) of average dF/F shows total GCaMP activity differs between the period of maximum respiratory depression and the onset of diaphragmatic rigidity (p = 0.026; paired t-test). Biii) Consistent with the temporal evolution of fentanyl, summary data (n=8) shows ventilation improves with the onset of rigidity (p = 0.0079). Improved ventilation also coincides with emergence of tonic diaphragm activation. Biv) Summary data of min ∫Dia EMG during the periods of respiratory depression and onset of rigidity. Ci) Representative trace of Z-Scored GCaMP activity (Top), plethysmography (middle), and minimum Diaphragm EMG under baseline and fentanyl. Shaded areas represent detected GCaMP burst events. Cii) Summary data of detected GCaMP burst events shows fentanyl stimulates LC bursting (p =0.023, paired t-test). Ciii) Ve inside LC GCaMP bursts was stimulated relative to outside LC GCaMP bursts under fentanyl but not baseline (p = 0.0003). Civ) However, min ∫Dia EMG activity inside LC GCaMP bursts was unchanged relative to outside LC GCaMP bursts in both conditions (p = 0.13). B-C: Paired t-test. Asterisk (*) indicates the difference between conditions;. One symbol = p < 0.05, two symbols = p < 0.01, three symbols = p < 0.001, four symbols = p < 0.0001.

### Oxygen sensitivity of diaphragmatic EMG activity during OIRD

Given OIRD is accompanied by discrete disruptions in oxygen homeostasis (Rai et al. 2025) and that the state of oxygenation can modulate the reversibility of fentanyl overdose (Haouzi et al., 2020), we next sought to determine how manipulating fractional inspired oxygen (FiO_2_) affected fentanyl-induced diaphragmatic rigidity. Neither hyperoxia (FiO_2_ = 100%) nor mild hypoxia (FiO_2 =_ 10%) alone induced expiratory ∫Dia EMG consistent with tonicity (data not shown). Moreover, under fentanyl, loss of phase coordinated (Inspiratory/Expiratory) diaphragm EMG activity was unaffected by hyperoxia (Fig 5Aii, p = 0.9) or mild hypoxia (Fig 5Bii, p = 0.9). On the other hand, severe hypoxia (FiO_2_ = 5%) shifted the fentanyl-induced breathing pattern into gasping, a stereotyped auto-resuscitative motor pattern (Gozal et al., 2002). This transition was characterized by elevated inspiratory diaphragm EMG activity (Fig. 6Ci, p = 0.0102). During gasping, tonic diaphragmatic EMG activity was abolished, expiratory EMG was reestablished to a level similar to baseline (Fig 6B, Cii, p = 0.23), and phasic EMG bursts were now locked to the inspiratory phase of each gasp and represented the dominant phase in EMG activity (Fig. 6Ciii, p = 0.52). Together, these experiments indicate that while fiDR is largely insensitive to changes in FiO_2_, fiDR is eliminated during gasping driven by severe-hypoxic challenge, suggesting that the fiDR may be the result of unique respiratory network states.

**Figure 5.**
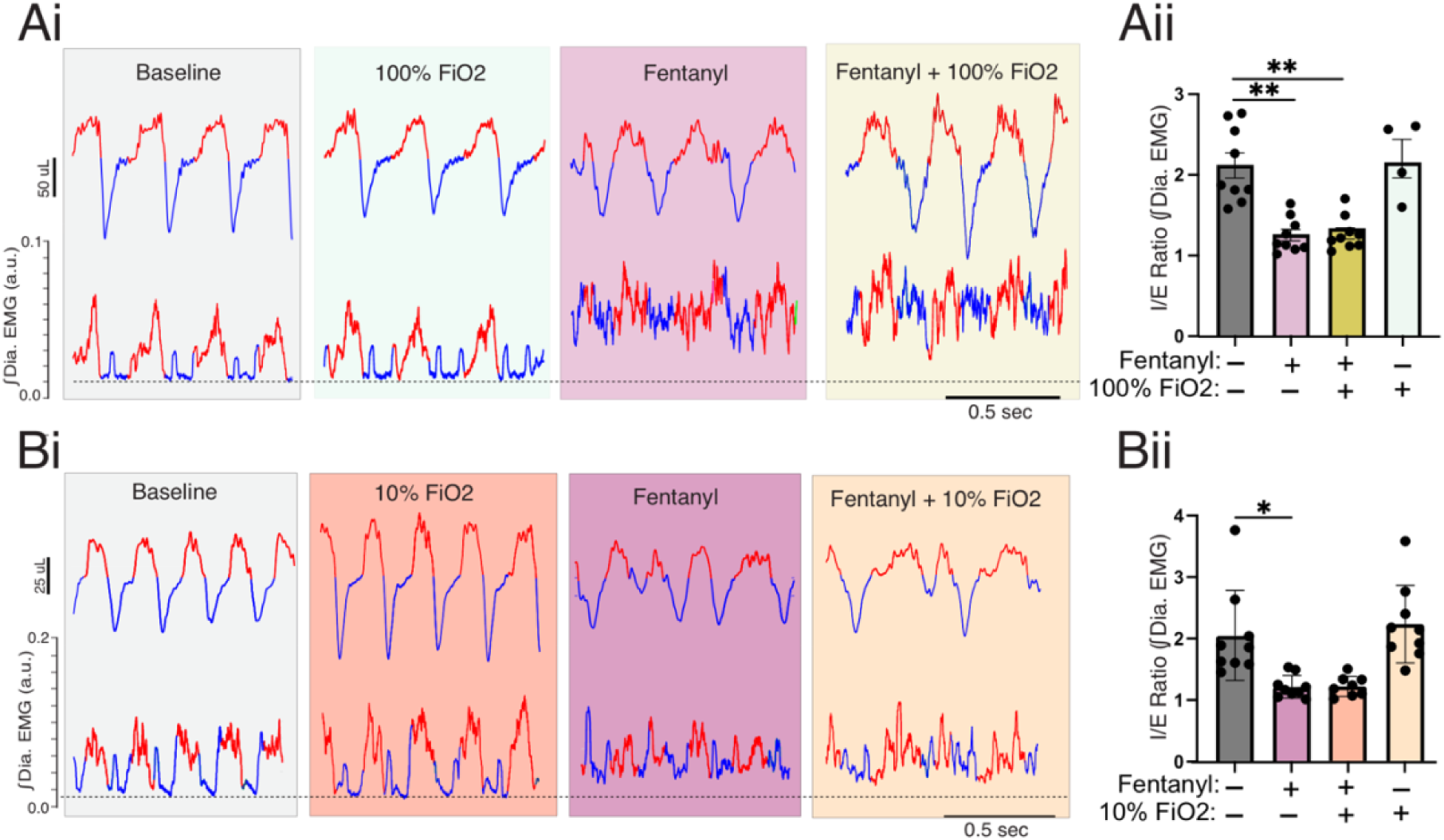
Fentanyl-induced diaphragmatic rigidity is unaffected under 100% -10% FiO_2_. Ai) Respiratory (Top) and integrated diaphragmatic EMG (Bottom) under Baseline, 100% FiO_2_, fentanyl, and fentanyl and 100% FiO_2_ with inspiration and expiration segmented. Aii) Summary data from the ratio of Inspiratory/Expiratory integrated diaphragmatic EMG activity under baseline, 100% FiO_2_, fentanyl, and fentanyl + 100% FiO_2_. Bi) Respiratory (Top) and integrated diaphragmatic EMG (Bottom) under Baseline, 10% FiO_2_, fentanyl, and fentanyl and 10% FiO_2_ with inspiration and expiration segmented. Bii) Summary data from the ratio of Inspiratory/Expiratory integrated diaphragmatic EMG activity under baseline, 10% FiO_2_, fentanyl, and fentanyl + 10% FiO_2_. Aii, Bii: Mixed Effects analysis with Tukey’s multiple comparisons. Asterisk (*) indicates the difference between conditions. One symbol = p < 0.05, two symbols = p < 0.01, three symbols = p < 0.001, four symbols = p < 0.0001.

**Figure 6.**
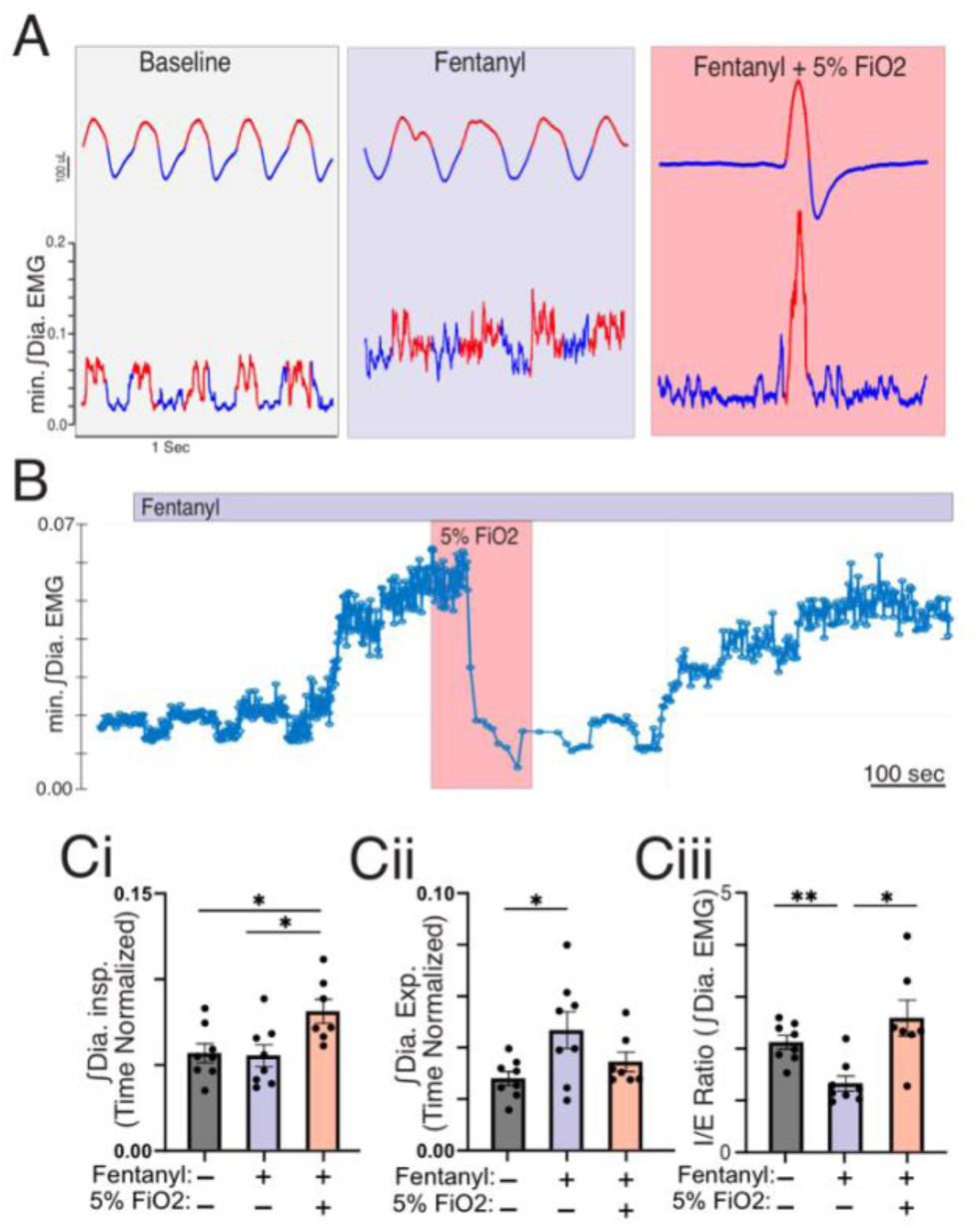
Gasping eliminates aberrant diaphragmatic activity induced by fentanyl. A) Respiratory (Top) and integrated diaphragmatic EMG (Bottom) under Baseline, 5% FiO_2_, fentanyl, and fentanyl + 5% FiO_2_ with inspiration (red) and expiration (blue) segmented. B) Trace of minimum integrated EMG activity shows fentanyl induced diaphragmatic tonicity that is reversed during gasping. C) Summary data (n = 7) of ∫Dia EMG shows under baseline (gray), fentanyl (purple), and fentanyl + 5% FiO_2_ (red) during gasping. Under fentanyl, severe hypoxia (5% FiO_2_) induced gasping increases inspiratory activity (Ci, p = 0.01) and rescues expiratory activity (Cii, p = 0.23) and inspiratory/expiratory ratio (Ciii, p = 0.52) to a level not different from baseline (One-way RM ANOVA with Tukey’s multiple comparisons). Asterisk (*) indicates the difference between conditions. One symbol = p < 0.05, two symbols = p < 0.01, three symbols = p < 0.001, four symbols = p < 0.0001.

The ability of 5% FiO_2_ to eliminate the fiDR raised the possibility that the phenomenon resulted from peripheral activation of the carotid bodies. Fentanyl exposure can augment carotid body (CB) activity during mild hypoxia (Peng et al., 2025) while near anoxic conditions can suppress CB activity (Peng et al., 2019). To test this possibility, we recorded diaphragmatic EMG in mice that had received bilateral CB denervation (CBD). Successful CBD was confirmed by the loss of the hypoxic ventilatory response to 10% FiO_2_ challenge (Fig 7B, p = 0.006). The degree of ventilatory depression induced by fentanyl was not significantly different in CBD mice from that of intact control animals (Fig 7B, p = 0.502). CBD mice did not develop tonic diaphragmatic activation under fentanyl (Fig 7A) as expiratory-phase EMG activity was reduced compared to SHAM controls (Fig. 7D, p = 0.002). However, bilateral diaphragmatic coordination was still significantly impaired in CBD mice under fentanyl (Fig. 7E, p = 0.009). Moreover, CB denervation did not prevent respiratory variability under fentanyl (Fig 7H, p = 0.037). These findings dissociate two components of fentanyl-induced diaphragmatic dysfunction: (1) tonic activation and expiratory-phase EMG elevation, which require intact CB afferent input; and (2) bilateral coordination failure, which is driven by central mechanisms independent of peripheral chemosensory input.

**Figure 7.**
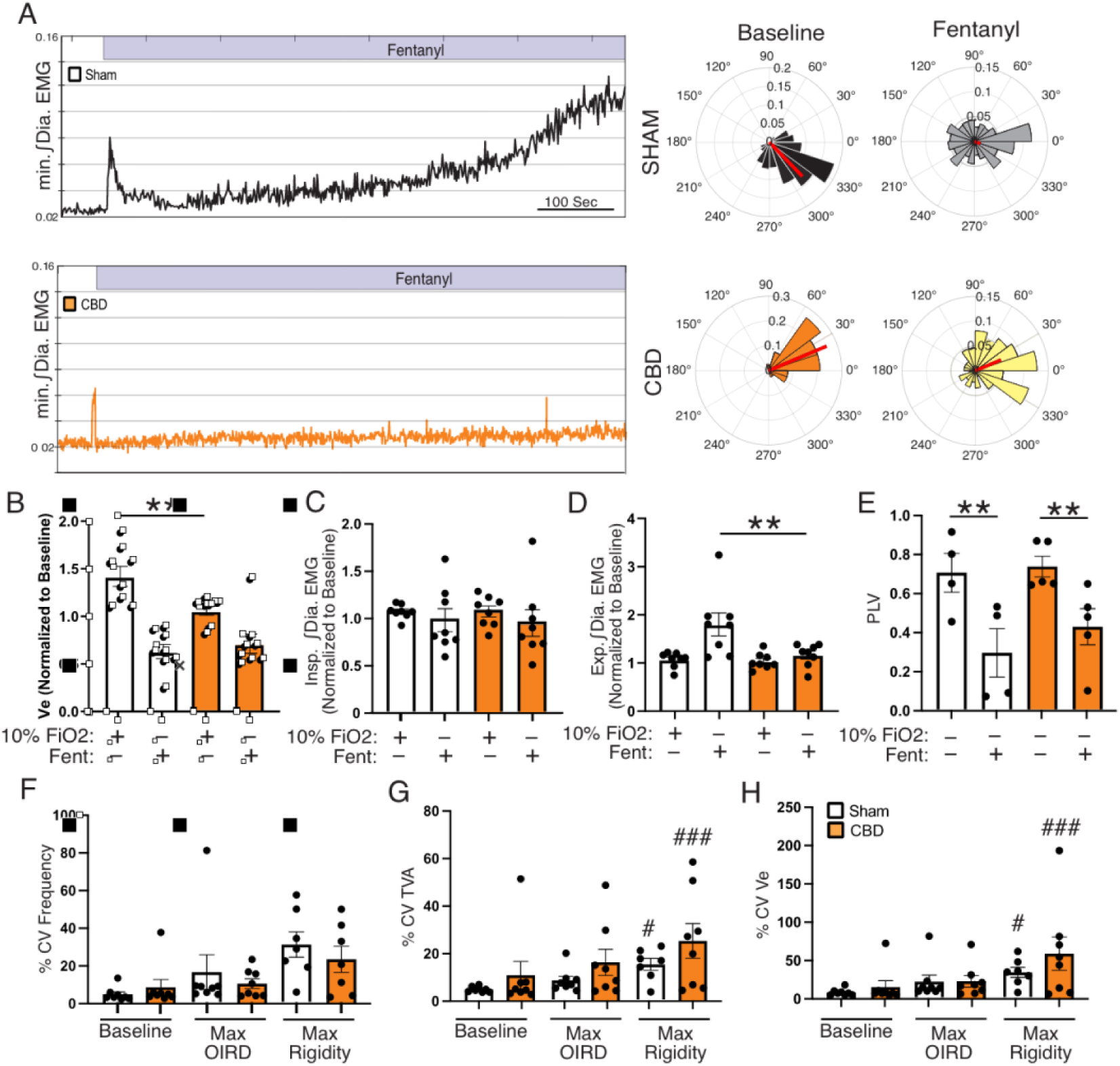
Carotid body denervation improves fentanyl-induced diaphragmatic tonicity. A) Left, Representative traces of minimum ∫Dia EMG activity from Sham (black) and Carotid body denervated (orange) animals. Right, polar plots of maximum Left-Right diaphragmatic EMG activity from SHAM and CBD mice under control and fentanyl conditions. B) Summary data of normalized Ve from Sham and CBD mice in response to 10% FiO_2_ and fentanyl shows loss of hypoxic ventilatory response (p = 0.006) with no change in OIRD (p = 0.50). C) Summary data of inspiratory ∫Dia EMG activity shows fentanyl had no effect on inspiratory activity in either SHAM or CBD mice. D) Summary data shows expiratory ∫Dia EMG activity was increased by fentanyl only in SHAM mice (p = 0.002). E) Despite CBD mice not exhibiting tonic diaphragm EMG under fentanyl, summary data of hemidiaphragm Phase-Lock values shows fentanyl reduces coordinated diaphragm activity in both SHAM (p = 0.004) and CBD (p = 0.009) mice. F-H) Summary data of coefficient of variance from Frequency (F), Tidal volume (G), and Ve (H) shows CBD mice exhibit respiratory discoordination similar to SHAM. B-E; Two-Way Anova with Tukey’s multiple comparison. F-H: Mixed effects with Tukey’s multiple comparison. Asterisk (*) indicates the difference between conditions, hashtag (#) indicated difference from baseline;. One symbol = p < 0.05, two symbols = p < 0.01, three symbols = p < 0.001, four symbols = p < 0.0001.

## DISCUSSION

In addition to suppressing ventilation, fentanyl can induce muscle rigidity. Fentanyl-induced muscle rigidity in respiratory muscles extends to the diaphragm, which can further aggravate overdose and complicate its treatment. Our study shows fentanyl-induced respiratory dysfunction as a temporally evolving syndrome, in which an initial phase of stable, centrally mediated ventilatory depression gives way to a later phase of unstable breathing accompanied by disrupted diaphragmatic activity. This disrupted activity is dissociated into generalized diaphragmatic tonicity and a previously undescribed loss of bilateral hemi-diaphragm coordination. While generalized tonicity involves peripheral input from the carotid bodies, loss of hemi-diaphragmatic coordination is independent of peripheral input. Our observations establish that ventilatory instability and bilateral motor discoordination are discrete, mechanistically separable features of overdose, and identify them as candidate targets for interventions aimed at the motor pathophysiology of overdose rather than respiratory depression alone.

We find diaphragmatic rigidity emerges after maximal ventilatory depression, and its onset coincided with partial recovery in Ve, accompanied by increased breath-to-breath variability. This temporal pattern suggests that such diaphragmatic rigidity is not the primary cause of the initial, most severe phase of ventilatory depression. While fentanyl initially depresses Ve in a manner consistent with MOR-mediated inhibition of central respiratory circuits, the subsequent emergence of tonic diaphragmatic activation, coupled with breath-to-breath instability, suggests a potential reorganization of respiratory motor output rather than a simple intensification of central depression. Such reorganization may result from distinct changes in the state of the respiratory network, in which respiratory instability persists despite improved Ve. Indeed, in humans, fentanyl exposure has been shown to produce ventilatory patterns consistent with Ataxic/Biot’s breathing (Farney et al., 2025; Wijdicks, 2007; Summ et al., 2022) that have the potential to further destabilize blood gas homeostasis and advance overdose pathophysiology in a feedforward manner. While these temporal dynamics are not captured by the classical definition of opioid-induced respiratory depression, our findings support and emphasize a multistage, progressive phenomenon in which respiratory discoordination emerges from, and outlasts, initial respiratory depression.

Diaphragmatic rigidity is driven by elevated activity during the expiratory phase, a feature that emerges during the later stages of fentanyl overdose. This phenomenon was not evident in adjacent expiratory musculature (Supplemental Figure 1), suggesting that the loss of fidelity in coordinated diaphragmatic activity does not simply reflect a generalized increase in muscle tone. Reorganization of the premotor respiratory control network to sustain respiratory motor output has been documented in several contexts, such as bilateral phrenic denervation in which animals continue to breathe despite loss of diaphragm function (LoMauro et al., 2021; Katagiri et al., 1994; Ahmed et al., 2012). Coordinated diaphragmatic contraction requires synchronous activation of bilateral phrenic motoneuron pools, which in turn depends on synchronized drive from the left and right preBötC (McKay and Feldman, 2008). Consistent with the reduction in phase-locked hemi-diaphragmatic activity, our slice experiments in the preBötC demonstrated that opioids perturb the bilateral burst-amplitude coupling between left and right preBötC networks. These observations support the hypothesis that fentanyl disrupts the network-level coupling responsible for motor coordination. Robust preBötC rhythmogenesis depends on a population of recruitable neurons whose synchronized activity underlies the generation of coordinated inspiratory output (Ashhad and Feldman, 2020). MOR activation can reduce network synchrony, thereby impeding but not eliminating the inspiratory rhythm. Moreover, reduced coordination in premotor drive may not be confined to preBotC as it projects bilaterally to premotor neurons of the rostral ventilatory group, which themselves communicate contralaterally before projecting to phrenic motor pools (Tan et al., 2010; Goshgarian et al., 1991). Thus, extensive bilateral connectivity within the respiratory network may increase vulnerability to an opioid-driven loss of bilateral synchrony as reduced premotor coordination at one level may be propagated and amplified across downstream levels of premotor control. Consistent with this interpretation, Olsen et al. (2026) recently showed that an opioid-like reduction in synaptic conductance among inspiratory populations alters burst timing, linking a breakdown in synaptic facilitation to discoordinated motor output. Fentanyl also causes a parallel disruption of phase coordination in vagal activity, where vagal firing becomes distributed across the respiratory cycle (Parks et al., 2025). Integrating these findings with our observations supports the view that the fentanyl-induced loss of phasic coordination is not limited to a single neuronal pathway but rather reflects a broader reorganization.

Aberrant diaphragmatic activity may arise from the interaction of opioid modulation and progressive blood gas disturbances during OIRD, whereby hypoxemia and/or hypercapnia amplify the opioid-mediated dysregulation of respiratory control. While increased variability in respiratory rate and tidal volume during the later phase of OIRD is consistent with ongoing homeostatic instability in response to fentanyl, manipulating inspired oxygen (FiO_2_) revealed that fentanyl-induced tonic diaphragm activity is not simply regulated by acute changes in arterial PO_2_. Neither hyperoxia (100% FiO_2_) nor modest hypoxia (10% FiO_2_) affected fentanyl-induced diaphragmatic dysfunction. In contrast, severe hypoxia (5% FiO_2_) following fentanyl administration triggered a well-characterized reconfiguration of breathing to gasping (Guntheroth and Kawabori, 1975, St-John and Paton 2004, Peña F. 2009) that was accompanied by an abolition of tonic diaphragmatic activity. The concurrent reduction in tonic diaphragmatic activity during gasping was reversible upon returning to a higher FiO_2_, suggesting that extreme hypoxia-driven gasping restores phasic diaphragm recruitment. As gasping reflects a unique network state in which activity is driven by medullary drive (Peña, 2009), the loss of pontine input (St-John and Paton, 2004), reconfiguration of the medullary networks (Potts and Paton, 2006), or a shift in neuromodulatory control of breathing (St-John and Leiter, 2008) drive the transition to a unique network state where phasic diaphragmatic activity is preserved despite the presence of fentanyl. Furthermore, we reasoned that aberrant diaphragmatic activity under severe hypoxia may also involve the withdrawal of tonic excitatory drive from CB chemosensory activity. While fentanyl can stimulate CB activity (Peng et al., 2025), CB output is suppressed during extreme hypoxia, when PO_2_ falls below the physiological range that drives chemosensory activity (Peng et al., 2019). CBD led to the abolition of tonic diaphragmatic activation and expiratory-phase EMG. However, CBD mice still exhibited hemi-diaphragmatic discoordination and ventilatory instability. Thus, we identified a previously undescribed role of CB activity in driving muscle rigidity and show that tonicity and discoordination are mechanistically distinct phenomena, each representing an independent pathophysiology during fentanyl overdose.

Bilateral coordination of preBötC bursting is vital for respiratory function, as complete loss of temporal burst coordination is lethal (Bouvier et al., 2010). The observed ventilatory instability during OIRD is similar to instability emerging with loss of coordinated left-right preBötC activity (Wu et al., 2017). In addition to showing that fentanyl induced hemi-diaphragmatic discoordination contributes to irregular and unstable breathing patterns during overdose, our findings support the idea that bilateral hemi-diaphragmatic discoordination involves lost left-right preBötC coupling independent of peripheral chemosensory input. In contrast, tonic diaphragmatic activation appears to require the continuous excitatory drive from CB afferents that fentanyl can produce.

CB drive is transmitted to the nucleus tractus solitarius and subsequently to premotor areas, including the LC, which may explain why generalized LC activity coincides temporally with the emergence of diaphragmatic rigidity. The unexpected finding that LC burst transients coincide with increased Ve but not with increased diaphragmatic rigidity also suggests changes in LC activity may be correlational rather than directly causing fentanyl-induced changes in diaphragmatic activity. However, as LC bursting consistently correlated with increased breathing during OIRD and general LC activity correlated with a rebound of Ve during fentanyl, we propose that the relationship between LC and ventilation may be contextually significant during overdose. Notably, the LC bursting did not stimulate breathing prior to fentanyl or following naloxone rescue, supporting the unique contextual feature of LC-ventilatory relationship and the potential importance of LC dynamics in shaping responses to fentanyl and its reversal. Rather than rejecting the model of LC driving fentanyl-induced muscle rigidity, our findings suggest that LC activation may modulate fentanyl-induced rigidity without being the sole determinant of diaphragmatic motor dysfunction. Furthermore, as LC is increasingly understood as a heterogeneous structure in which distinct projection modules serve distinct functions (Ma et al., 2023), it remains possible that a specific subset of LC neurons drives diaphragmatic tonicity even when aggregate population activity does not predict it.

Beyond reframing the conceptual model of OIRD, our findings extend past the canonical view of overdose as central respiratory depression. Current clinical understanding of OIRD remains centered on the suppression of respiratory drive, while the broader motor pathophysiology of overdose is comparatively undefined (Boom et al., 2012; Bateman et al., 2023). The contribution of diaphragmatic rigidity to ventilation, and the respiratory variability that accompanies it, have not been well characterized. We propose that respiratory variability is an underappreciated feature of fentanyl overdose. The breath-to-breath instability we observe is reminiscent of the ataxic breathing associated with opioid exposure (Farney et al. 2025,Walker et al., 2007), a pattern that is predicted to destabilize arterial blood-gas homeostasis. This scenario is further complicated because opioids perturb metabolism through several mechanisms (i.e. altering oxygen consumption, thermogenesis, and metabolic rate) (Romanovsky and Blatteis, 1996; Romanovsky and Blatteis, 1998), and driving changes in tissue oxygenation that arise from both respiratory depression and vascular tone (Solis et al., 2017). Species differences are an important consideration. Mice are comparatively opioid-tolerant, and exhibit pronounced strain-dependent variation in opioid lethality (Marchette et al., 2023), raising the possibility that variability is relatively well tolerated in this species. Whereas, in more susceptible species, the same instability could precipitate respiratory collapse. Given complete loss of left–right preBötC coordination can be lethal (Bouvier et al., 2010), the bilateral premotor discoordination that we describe here could be exaggerated in species with lower opioid tolerance. Lastly, we find that not all subjects exhibited disrupted diaphragmatic activity in response to fentanyl. This observation is consistent with previously described variability observed in opioid susceptibility and tolerance development (Galer et al., 1992; Solhaug and Molden, 2017; Kest et al., 2002). Specifically, we have previously described divergent respiratory endotypes to development of OIRD tolerance such that, while most animals develop tolerance following repeat opioid use, tolerance fails to develop in a minority of mice (Rai et al. 2025).

Collectively, our work challenges the prevailing framework of OIRD as a single phenomenon of central respiratory depression. Rather, we propose fentanyl overdose is a syndromic phenomenon that evolves with the time course of overdose. In adult mice, we find that the initial phase of fentanyl overdose is characterized by stable, centrally mediated depression. This stable depression gives way to a later phase of unstable breathing that coincides with perturbed diaphragmatic activity. This perturbed activity appears to involve distinct roles for peripheral chemosensory activity and a loss of premotor coordination. Thus, our study dissociates the central and peripheral contributions to the effects of fentanyl on respiratory motor output and emphasizes the need to reconsider the role of coordinated drive and peripheral contribution to effectively combat fentanyl overdose.

## AUTHOR CONTRIBUTIONS

JSP and AJG conceived and designed experiments; JSP, GEF, BB, and SWW performed experiments; YHF performed CB denervation; JSP, JF, GEF, BB and SWW conducted analyses; JSP and AJG drafted the manuscript; All authors revised the manuscript. WS and AJG provided reagents and support.

---

**Supplemental Figure 1.**
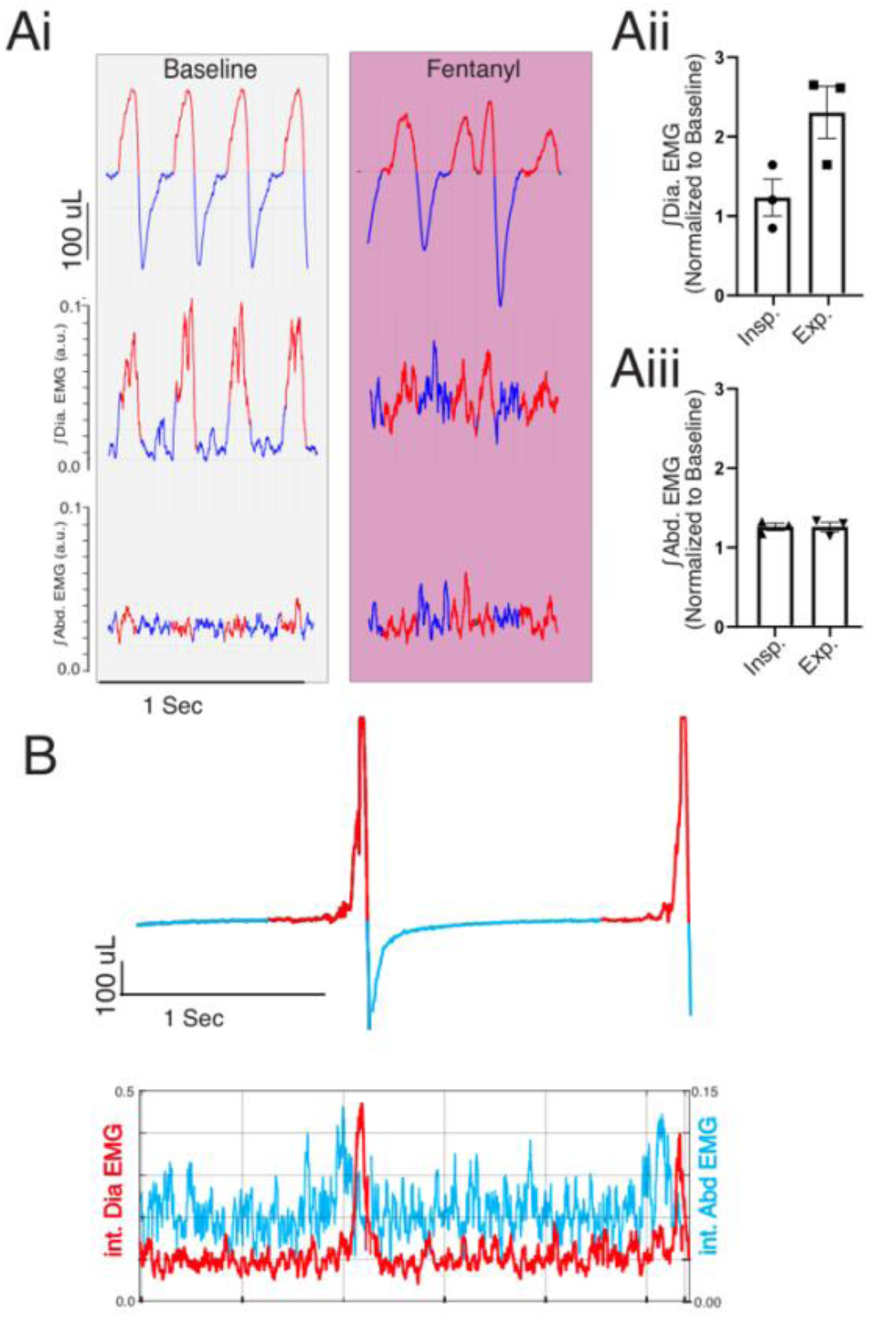
Aberrant diaphragmatic activity induced by fentanyl is independent of abdominal activation. Ai) Representative traces of respiration (Top), integrated diaphragm (middle), and integrated abdominal (bottom) EMG activity. Red indicated inspiration and blue indicates expiration. Aii) Summary data of integrated diaphragm activity under fentanyl (normalized to baseline) in the inspiratory and expiratory phases. Aiii) Summary data of integrated abdominal activity under fentanyl (normalized to baseline) in the inspiratory and expiratory phases. B) Representative trace of breathing (top) and ∫EMG (bottom) activity from the diaphragm (red) and abdominal (blue) during 5% FiO_2_ induced gasping. Aii-iii: paired t-test. Asterisk (*) indicates the difference between conditions, hashtag (#) indicated difference from baseline. One symbol = p < 0.05, two symbols = p < 0.01, three symbols = p < 0.001, four symbols = p < 0.0001.

**Supplemental Figure 2.**
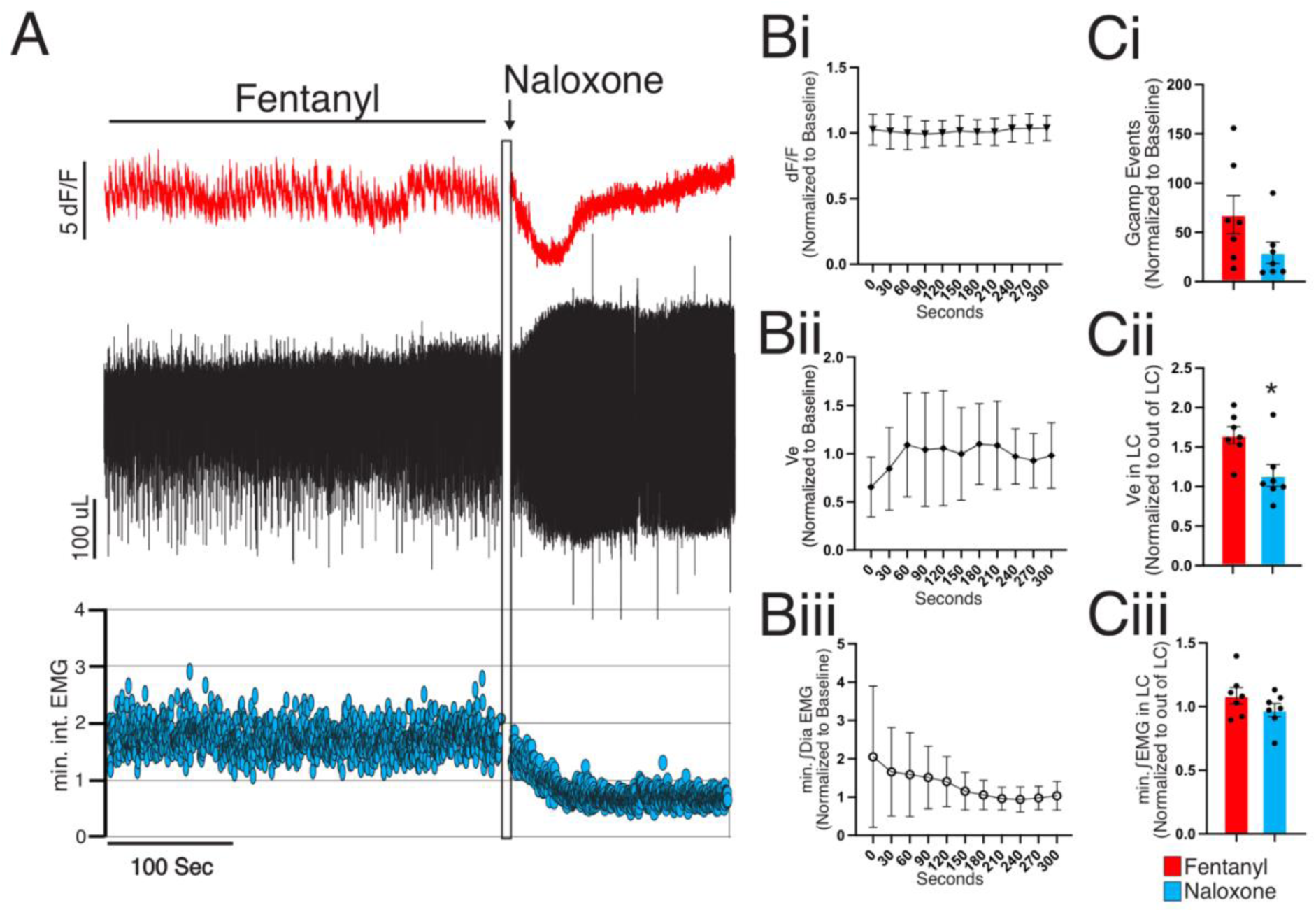
LC coupled ventilatory bursts are reversed by naloxone. A) Representative trace of dF/F (Top), plethysmography (middle), and minimum Diaphragm EMG (bottom) under fentanyl (i.p. 0.7mg/kg) and naloxone (i.p. 2mg/kg). Bi-iii) Time course of normalized LC dF/F (Bi), Ve (Bii), and minimum òDiaphragm EMG (Biii) under fentanyl and following reversal with naloxone (indicated by arrow). Ci) Summary data of detected GCaMP burst under fentanyl and following naloxone (p = 0.096). Cii) Fentanyl induced Ve stimulation inside LC GCaMP bursts relative to outside was eliminated following naloxone (p = 0.038). Ciii) min ∫Dia EMG activity inside LC GCaMP bursts was unchanged relative to outside LC GCaMP bursts following naloxone reversal (p = 0.15). Ci-iii: paired t-test. One symbol = p < 0.05, two symbols = p < 0.01, three symbols = p < 0.001, four symbols = p < 0.0001.

